# Chemical conversion of human conventional Pluripotent Stem Cells to Trophoblast Stem Cells

**DOI:** 10.1101/2022.04.07.487453

**Authors:** Irene Zorzan, Riccardo Massimiliano Betto, Giada Rossignoli, Mattia Arboit, Andrea Drusin, Paolo Martini, Graziano Martello

**Author notes:** These authors contributed equally (Irene Zorzan, Riccardo Massimiliano Betto).

## Abstract

In human embryos, naive pluripotent cells of the inner cell mass generate epiblast, primitive endoderm and Trophectoderm (TE) lineage, whence trophoblast cells derive. *In vitro*, naive pluripotent stem cells (PSCs) retain this potential and can generate trophoblast stem cells (TSCs), while conventional PSCs form amnion-like cells and lack the competence to generate TSCs. Transient histone deacetylase and MEK inhibitions with LIF stimulation can be used to chemically reset conventional to naive PSCs. Here we report that chemical resetting induced expression of both naive and TSC markers and of placental imprinted genes. A modified chemical resetting protocol allowed for the fast and efficient conversion of conventional PSCs into TSCs, entailing shutdown of pluripotency genes and full activation of the trophoblast master regulators, without induction of amnion markers. Chemical resetting generates a responsive intermediate state, in which conventional PSCs rapidly acquire competence to form TSCs without the need of stabilisation and expansion in a naive state. The efficiency and rapidity of our system will be useful for the study of cell fate transitions, and to generate models of placental disorders.

## Introduction

The placenta is composed of cells of both maternal and embryonic origin, the latter known as trophoblast. Cytotrophoblasts (CTB) are highly proliferative cells that, in turn, give rise to syncytiotrophoblast (STB) and extravillous cytotrophoblast^1,2^. CTBs have been recently captured *in vitro* as trophoblast stem cells (TSCs)^3^ from first-trimester placental tissues, using a medium (TSC medium) containing epidermal growth factor (EGF), a GSK3 inhibitor (CHIR99021), two inhibitors of transforming growth factor beta (A83-01 and SB431542), an inhibitor of histone deacetylase (Valproic Acid, VPA), and an inhibitor of Rho-associated kinase (Y27632)^3^.

In the early blastocyst stage embryo, two cell populations are generated as the result of the first cell fate decision, the ICM and the TE. The TE will generate CTBs after implantation^3–5^, while the ICM gives rise to all somatic cells. The ICM of human embryos is more plastic than its murine counterpart, as it is able to generate TE^6^, as suggested by single-cell RNA-sequencing analyses of early embryos^7^ and recently confirmed by *in vitro* differentiation of explanted ICM^6^.

Human naive PSCs have been derived either directly from pluripotent cells of the preimplantation embryo^8,9^, from resetting of conventional PSCs by transgene expression^8,10^ or using a combination of inhibitors and cytokines^8,11,12^ or by reprogramming from human fibroblasts^13–16^.

Human naive PSCs retain *in vitro* the plasticity of the ICM and can readily differentiate towards TE^4,6^ and TSCs^11,17–19^. Several different combinations and cytokines have been used for the expansion of naive PSCs^8,10,11,20,21^. Among them we chose the PXGL medium^12^, containing the cytokine LIF, inhibitors of the MEK and PKC kinases and of tankyrase.

Conventional PSCs are in a pluripotent state primed for differentiation, more akin to the post-implantation epiblast^22–24^. When exposed to TSC medium, conventional PSCs fail to form TSCs and acquire a neuroectodermal fate^17^ or stop proliferating and experience elevated cell death^19^.

Several studies reported the formation of trophoblast-like cells from conventional PSCs after BMP stimulation^5,25–29^. The identity of those trophoblast-like cells has been put into question^26^, as they do not fulfil established criteria for trophoblast identity^5^ and recent reports showed that they actually correspond to amnion^4,6^. Indeed, BMP is not involved in trophectoderm specification in the human blastocyst^30^ and conventional PSCs have recently been reported to differentiate into amnion-like cells in response to BMP^31^. All these studies indicate that conventional PSCs can generate amnion-like cells and lack the competence for trophoblast differentiation.

Whether TSC differentiation competence can be reinstated only by full acquisition of naive pluripotency is still an open question, but several reprogramming studies would indicate otherwise. Single-cell analyses of cells acquiring naive pluripotency revealed the existence of multiple trajectories, depending on the experimental conditions used. In the case of resetting of murine primed PSCs, cells either follow a direct trajectory or go via developmental states slightly more advanced (mesoderm) or more primitive (ICM), eventually all converging to the naive pluripotent state^32^. In the case of somatic cell reprogramming achieved by a combination of chemicals or using the GETMS transcription factor cocktail, cells first transiently acquire an extraembryonic fate (i.e. primitive endoderm) before reaching and stabilising in the naive pluripotent state^33,34^. Furthermore, different cell populations might be generated, as in the case of trophoblast-like cells generated along with pluripotent cells, during reprogramming of human somatic cells^35^. Those trophoblast-like cells were directly captured as TSCs, without the need of prior stabilisation or expansion in a naive pluripotent state. Overall, these studies suggest that some resetting and reprogramming protocols confer competence for generation of both pluripotent and extraembryonic cells.

In this study we observed that chemical resetting of human conventional PSCs generated a mixed cell population, containing both pluripotent and non-pluripotent cells. Based on the activation of placental imprinted transcripts and markers of TE and TSC, we hypothesised that the mixed cell population could contain trophoblast-like cells, which we could capture and expand as TSCs. We were indeed able to convert conventional PSCs to TSCs in an efficient and rapid manner, with no need of intermediate expansion under conditions supporting naive pluripotency.

## Results

### Activation of TE and trophoblast markers upon chemical resetting of conventional PSCs

Human conventional induced pluripotent stem cells (iPSCs) obtained from keratinocytes (KiPS)^10^ were reset to a naive state by transient histone deacetylase inhibition^12^ (Fig. 1a), a process described as chemical resetting. After 7 days, several dome shaped, compact colonies were present, that we could readily expand for multiple passages in PXGL medium^36^ (Fig. 1b, green arrows). These colonies were composed of cells expressing the naive pluripotency marker KLF17 and the general pluripotency factor POU5F1/OCT4 (Fig. 1c).

**FIGURE 1.**
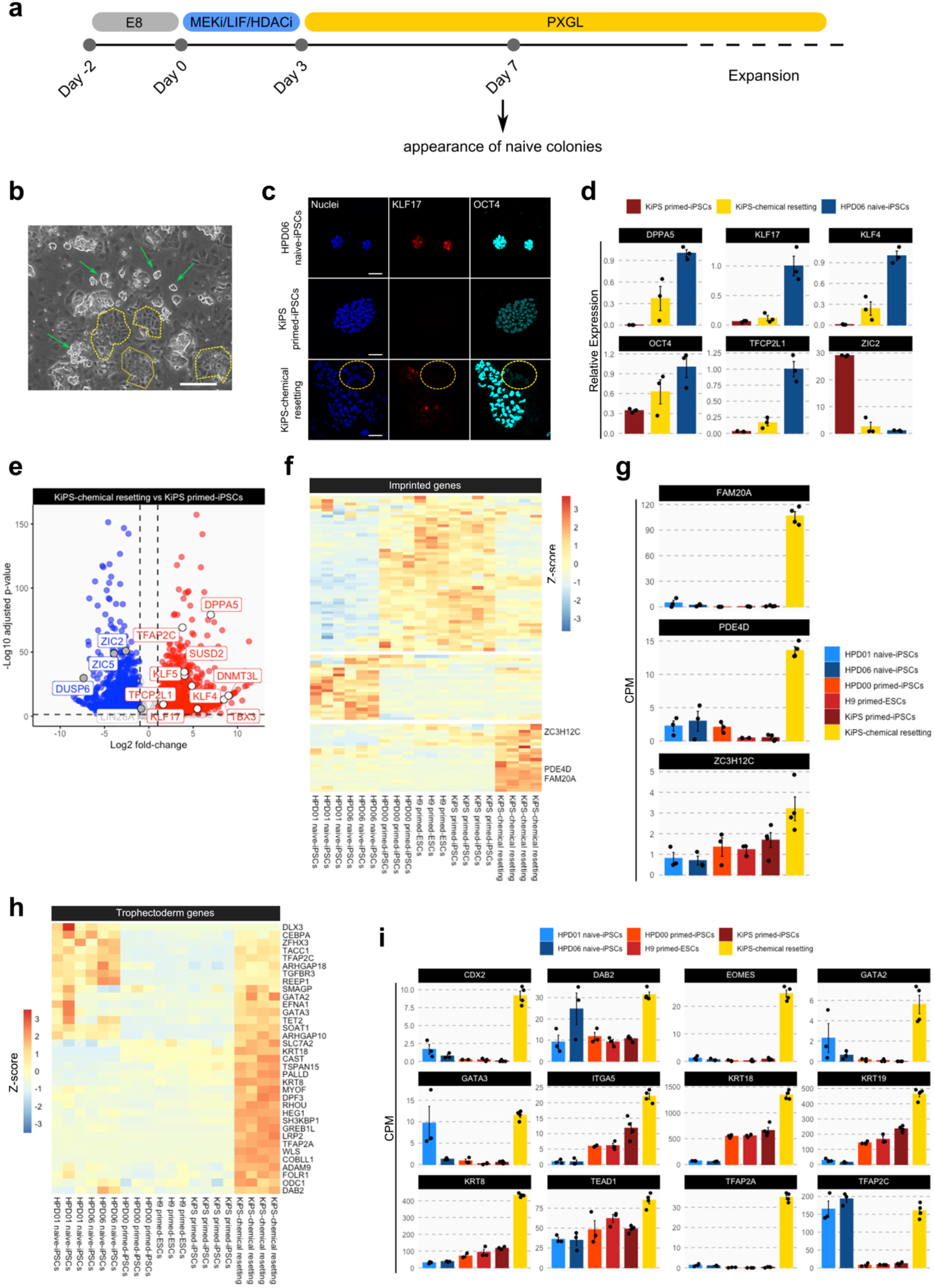
Activation of TE and trophoblast markers upon chemical resetting of conventional PSCs. **a** Experimental strategy for chemical resetting of human primed PSCs to naive pluripotency. **b** Morphology of KiPS-chemical resetting after 14 days. Green arrows indicate naive colonies, dashed yellow lines indicate clusters of flat polygonal cells. Representative images of three independent experiments are shown. Scale bar: 200 μm. **c** Immunostaining for the naive pluripotency marker KLF17 and the shared pluripotency marker POU5F1/OCT4. Representative images of three independent experiments are shown. Scale bar: 50μm **d** Gene-expression analysis by qPCR of KiPS primed-iPSCs, KiPS-chemical resetting and HPD06 naive-iPSCs. Bars indicate the mean ± SEM (standard error of the mean) of three independent experiments shown as dots. Expression was normalised to the mean of HPD06 naive-iPSCs samples. **e** Transcriptome analysis of KiPS-chemical resetting compared to KiPS primed-iPSCs. DOWN-regulated (Log2 fold-change < -1 and adjusted p-value < 0.05) and UP-regulated (Log2 fold-change > 1 and adjusted p-value < 0.05) genes are indicated in blue and red respectively. Known primed pluripotency and naive pluripotency markers are highlighted as grey and white dots respectively. Adjusted p-values were calculated with Wald Test. **f** Heatmap of 93 imprinted genes detected in pluripotent cells. Z-Scores of row-scaled expression values (CPM) are shown. Red and blue indicate high and low expression, respectively. Three main clusters have been identified, imprinted genes highly expressed either in primed PSCs, in naive PSCs or after chemical resetting, respectively. Three placental imprinted genes (AM20A, PDE4D and ZC3H12C) are highlighted in the latter cluster. **g** Barplots showing the absolute expression (CPM) of three placental imprinted genes in the reported conditions highlighted in different colours. Bars indicate the mean ± SEM of independent experiments shown as dots. **h** Heatmap of trophoblast specific genes^7,40^ UP-regulated (Log2 fold-change > 1 and adjusted p-value < 0.05) in KiPS-chemical resetting vs KiPS primed-iPSCs comparison. Z-Scores of row-scaled expression values (CPM) are shown. Red and blue indicate high and low expression, respectively. **i** Barplots showing the absolute expression (CPM) of twelve trophoblast genes found to be significantly UP-regulated in KiPS-chemical resetting vs KiPS primed-iPSCs comparison. Bars indicate the mean ± SEM of 3 independent experiments shown as dots.

As previously reported^12^, the cell population obtained after chemical resetting was heterogeneous (Fig. 1b) and also contained flat polygonal cells (Fig. 1b, dashed yellow lines) in addition to compact, naive-like colonies. These cells also did not express pluripotency markers (Fig. 1c, yellow circle). We asked ourselves what those cells might be.

We performed quantitative Polymerase Chain Reaction (qPCR) on the mixed culture obtained and observed partial activation of the naive-specific markers DPPA5, KLF17, KLF4 and TFCP2L1 (Fig. 1d), consistent with the fact that only a fraction of cells display naive features. ZIC2, a marker specifically expressed by conventional PSCs, was greatly reduced after chemical resetting (Fig. 1d). We performed RNA-sequencing (RNA-seq) and confirmed the activation of naive-specific markers (DPPA5, KLF4/5/17, TFCP2L1 and SUSD2) and the downregulation of markers of conventional PSCs (ZIC2/5, DUSP6, LIN28A - Fig. 1e). We concluded that the chemical resetting generates a mixed culture of naive PSCs and additional undefined cell types, which are unlikely to be conventional PSCs. Imprinted genes are expressed in a stage-specific and tissue-specific manner^37^. We interrogated RNA-seq data and observed that naive and conventional PSCs express very distinct sets of imprinted genes (Fig. 1f). Strikingly, we observed a group of imprinted genes highly expressed only after chemical resetting. Among them we found placental transcripts, such as FAM20A, PDE4D and ZC3H12C^38^ (Fig. 1g). Placental imprinted genes are highly expressed in the trophoblast cells in the placenta^39^, so we asked whether other markers of the TE lineage were induced after chemical resetting. We observed that 35 out of 107 expressed genes (p-value=4.27 × 10^−6^, Fisher’s exact test) specifically expressed by TE cells in the human embryo^40^ were highly induced after chemical resetting (Fig. 1h), including KRT19, DAB2, LRP2 and TFAP2A (Fig. 1h). We also analysed a wide range of markers expressed by TSCs *in vitro* and observed strong upregulation after resetting (Fig. 1i). Of note, human naive PSCs display spontaneous and low expression levels of some trophoblast markers, consistently with their propensity to generate TSCs^17^. However, the levels observed after chemical resetting were significantly higher than those of both primed and naive PSCs (Fig. 1i). Interestingly, TFAP2C, a gene required for self-renewal of both naive PSCs and TCSs^6,41^, was fully activated after chemical resetting.

Our data indicate that chemical resetting leads to activation of TE and TSC genes.

### TSCs are generated via chemical resetting of conventional PSCs

Trophoblast stem cells (TSCs) are stably expandable cell lines recently obtained from early human embryos^3^ or via differentiation of human naive PSCs^6,17,18^. Conventional PSCs are unable to generate *bona fine* TCSs^17,19^, as they are too developmentally advanced and lost the competence to do so. We could readily obtain TSCs from naive PSCs (both human embryonic stem cells and iPSCs^10,13^) and validated their correct identity by the expression of TSC markers GATA2/3, KRT7 and TFAP2A and by their capacity to differentiate towards STBs (Fig. 2a-b) and to form trophoblast organoids^42^ actively producing human chorionic gonadotropin (Fig. 2c-d). RNA-seq data further confirmed a distinct global expression profile of these naive PSCs-derived TSC lines compared to naive and conventional PSCs (Fig. 2e). These TSC lines, named H9 naive-TSCs and HPD06 naive-TSCs, served as positive controls in the following experiments.

**FIGURE 2.**
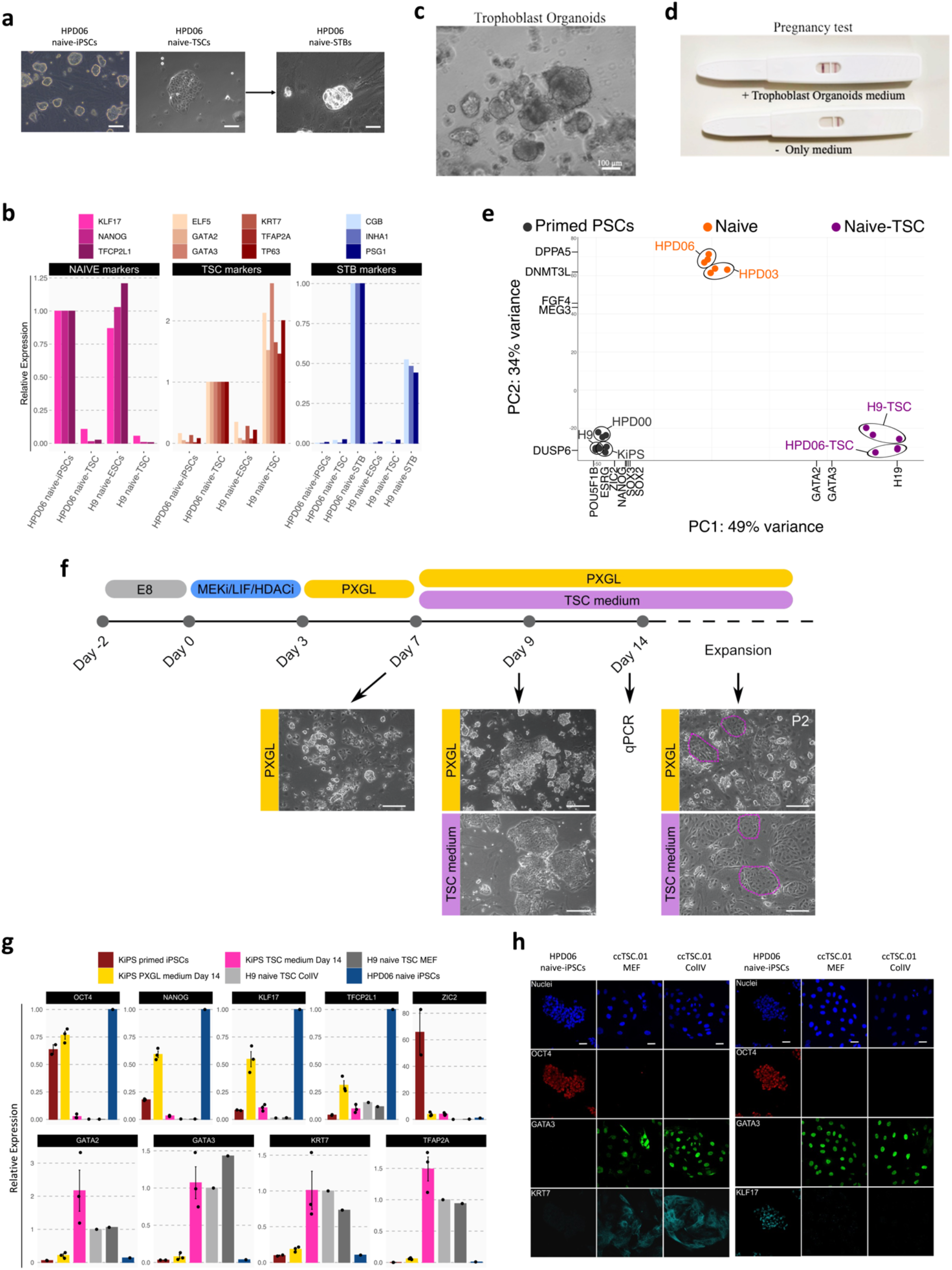
TSCs are generated via chemical resetting of conventional PSCs. **a** Phase contrast images of naive hPSC (HPD06), hTSC cells derived from naive hPSCs (HPD06 naive-TSCs) and syncytium trophoblast (STB) cells derived from HPD06 naive-TSCs. Similar results were obtained with H9 naive-ESCs and HDP03 naive-iPSCs. Scale bars: 400 μm. **b** Gene expression analysis by RT–qPCR for naive (KLF17,TFCP2L1 and NANOG) and trophoblast (GATA3, GATA2, KRT7, ELF5, TP63 and TFAP2A) marker genes in naive PSCs and hTSC cells, and for STB marker genes (CGB, INHA1 and PSG1) in naive PSCs, TSC cells and STB derived from two different naive PSC lines (HPD06 and H9). **c** Phase-contrast images of trophoblast organoids derived from TSCs plated in Matrigel drop and exposed to trophoblast organoids medium (TOM^42^) for 10 days. Scale bars: 100 μm. **d** Pregnancy test detecting human chorionic gonadotropin in trophoblast organoid medium. **e** Principal component analysis of primed-iPSCs (HPD00, H9 and KiPS), naive-iPSCs (HPD06 and HPD01) and TSC cells derived from naive PSCs (HPD06 naive-TSCs and H9 naive-TSCs) performed on the top 5000 most variable genes identified through RNA-seq. Conditions are highlighted in different colours and biological replicates of the same cell line are labelled. **f** Experimental scheme of the conversion of conventional PSCs into naive PSCs or TSCs. Morphologies of cells at day 7 in PXGL medium, at day 9 in PXGL medium or in TSC medium, and after 2 passages in the two culture conditions are shown. The TSCs obtained could be readily expanded and were named ccTSC.01. Representative images of 2 biological replicates are shown. Scale bars: 200 μm. **g** Gene-expression analysis by qPCR of KiPS primed-iPSCs, KiPS in PXGL medium at day 14, KiPS in TSC medium at day 14, H9 naive-TSC on ColIV, H9 naive-TSC on MEF and HPD06 naive-iPSCs. Top: naive, primed and general pluripotency markers, expression was normalised to HPD06 naive iPSCs; bottom: trophoblast markers, expression was normalised to H9 naive TSC. Bars indicate the mean ± SEM of biological replicates shown as dots. **h** Immunostaining for the pluripotency markers OCT4 and KLF17, and the TSC markers GATA3 and KRT7 of HPD06 naive-iPSC and ccTSC.01 culture both on MEF and on collagen IV (ColIV) after 3 passages. Representative images of 2 biological replicates are shown. Scale bars: 30 μm.

Based on the strong activation of TSC genes (Figure 1) upon chemical resetting, we hypothesised that chemical resetting might endow conventional PSCs with the competence to generate TSCs. To test this hypothesis, we modified the chemical resetting protocol and, after 7 days of resetting, cells were exposed to either naive PSC- or TSC-supporting conditions (Fig. 2f). As expected, several dome shaped colonies emerged in PXGL (Fig. 2f), along with flat polygonal cells whose morphology resembles TSCs (purple dashed lines in Fig. 2f). Transcriptional analysis confirmed upregulation of naive specific and general pluripotency markers (Fig. 2g).

Strikingly, under TSC conditions we observed complete repression of pluripotency markers and activation of TSC markers to levels observed in established TSCs (Fig. 2g). The cells obtained, which we named ccTSCs (chemically converted TSCs, ccTSC.01) expanded robustly for several passages, with a stable and very homogeneous morphology. Immunostaining confirmed the presence of TSC markers GATA3 and KRT7, and complete absence of pluripotency markers OCT4 and KLF17 (Fig. 2h). TSCs have been expanded both on a feeder-layer of mouse embryonic fibroblasts (MEFs) or on Collagen IV (ColIV) coated wells^3,4,6,19^. Once established, we expanded ccTSCs for multiple passages under both conditions with homogenous expression of TSC markers and similar expansion kinetics (Fig. 2h). We conclude that a modified chemical resetting protocol allowed the efficient conversion of conventional PSCs to TSCs.

### Rapid and highly-efficient generation of ccTSCs

We obtained ccTSCs after 3 days of chemical resetting, followed by 4 days in PXGL, a medium supporting human naive PSCs (Fig. 2f-h). We asked whether exposing cells directly to TSC medium right after 3 days of chemical resetting would also lead to ccTSC generation, skipping an intermediate step in PXGL (Fig. 3a).

**FIGURE 3.**
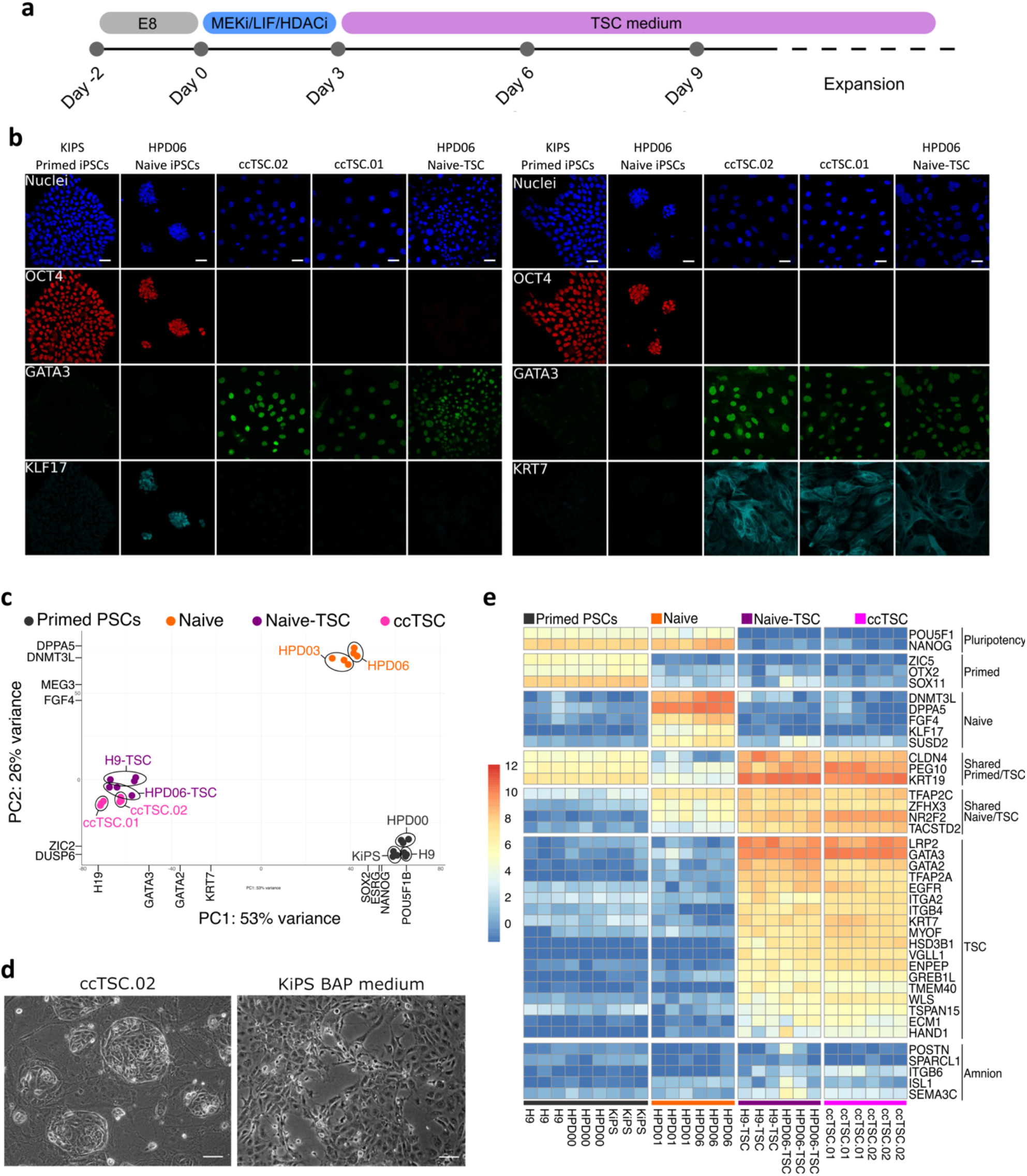
Rapid and highly-efficient generation of ccTSCs. **a** Experimental scheme of the conversion of primed PSCs into ccTSCs. **b** Immunostaining for the pluripotency markers OCT4 and KLF17, and the TSC markers GATA3 and KRT7 of KiPS primed-iPSCs, HPD06 naive-iPSCs, ccTSC.02 at passage 4, ccTSC.01 at passage 4 and HPD06 naive-TSC. Scale bars: 30 μm. **c** Principal component analysis of ccTSCs (ccTSC.01 and ccTSC.02) with primed-PSC (HPD00, H9 and KiPS), naive-iPSCs (HPD06 and HPD01), and TSCs derived from naive PSCs (HPD06 naive-TSCs and H9 naive-TSCs) performed on the top 5000 most variable genes identified through RNA-seq. Conditions are highlighted in different colours and biological replicates of the same cell line are labelled. **d** Phase-contrast images of ccTSC.02 at passage 5 and of BAP-treated KiPS after 3 days of treatment. Scale bar: 200 µm. **e** Heatmap of primed, naive and trophoblast specific genes in primed-PSCs (HPD00, H9 and KiPS), naive-iPSCs (HPD06 and HPD01), TSCs derived from naive PSCs (HPD06 naive-TSCs and H9 naive-TSCs), and ccTSCs (ccTSC.01 and ccTSC.02). Row-scaled log2(CPM) expression values are shown. Red and blue indicate high and low expression, respectively. Main clusters have been identified as marker genes for general pluripotency, primed-iPSCs, naive-iPSCs, TSCs and shared markers between groups^17^, as well as amnion markers^4^.

Much to our surprise, after only 3 days in TSC medium, we detected full activation of TSC markers and repression of markers of both naive and conventional pluripotency (Supplementary Fig. 1). Accordingly, the entire cell population (ccTSC.02; Fig. 3d, left panel) displayed a morphology indistinguishable from TSCs obtained from naive PSCs (Fig. 2a). The ccTSCs could be readily expanded and displayed robust and homogenous expression of TSC markers and absence of pluripotency markers (Fig. 3b). Moreover, global transcriptome analysis revealed that both ccTSC lines clustered with H9 naive-TSCs and HPD06 naive-TSCs (Fig. 3c). We conclude that a transient inhibition of histone deacetylases is sufficient to allow efficient and rapid conversion of conventional PSCs to ccTSCs, without need for an intermediate step in a naive PSC-supporting medium.

Exposure of conventional PSCs to Bone Morphogenetic Proteins (BMPs), either alone or in combinations with the ALK4/5/7 inhibitor A83-01, and the FGFR inhibitor PD173074 (BAP medium) has been used to generate cells with some molecular features of trophoblast cells^25,43^. However, several groups reported that BAP cells differ from primary trophoblast cells^26^ and from *bona fide* TCS^4,6,17^. BAP cells do not fulfil stringent criteria for trophoblast identity^5,26^ and, rather, display transcriptomes consistent with amnion identity. Although we have not used BMP ligands in our protocol, we decided to investigate whether our ccTSC were distinct from BAP cells. We exposed conventional KiPS cells to BAP medium and observed a cell morphology consistent with previous reports^4,43^ and completely distinct from our ccTSCS (Fig. 3d). Gene expression analysis confirmed activation of KRT7 and GATA3, but also of amnion markers IGFBP3^4,6^ (Supplementary Fig. 1b) In contrast, HAVCR1, a marker of *bona fide* TSCs^4,6^, was not induced in BAP cells (Supplementary Fig. 1b). In stark contrast, ccTSCs showed robust induction of several TSC markers and negligible expression of amnion markers, further indicating their *bona fide* TSC identity (Fig. 3e). Interestingly, some TSC markers, such as TFAP2C and TACSTD2, were found to be expressed at low levels in naive-iPSCs lines too, in line with recent findings of co-expression of embryonic and extraembryonic genes in naive-iPSCs as a distinctive feature of naive pluripotency^17^ (Fig. 3e). However, we also found expression of TSC markers (i.e. CLDN4, PEG10 and KRT19) in KiPS primed-iPSC, even if at lower level (Fig. 3e).

In sum, conventional PSCs were rapidly and efficiently converted into ccTSCs, which display molecular features compatible with *bona fide* TSCs.

Transient histone deacetylase inhibition followed by exposure to PXGL is applied for routine generation of human naïve PSCs from conventional PSCs^12^. Using the same procedure, we could rapidly obtain TSCs through the exposure of conventional PSCs to TSC medium right after 3 days of chemical resetting (Fig. 3). We therefore wondered if the KiPS primed-iPSCs to TSCs conversion proceeds through an intermediate naive-like pluripotent state with transient naive markers activation or directly to TSCs. We performed RNA-seq analysis at different timepoints during the histone deacetylase inhibition step and subsequent stabilisation in PXGL or TSC medium (Fig. 4a). During the initial 3 days of chemical resetting, we observed upregulation of markers of naive pluripotency and of TSCs (Fig. 4b). At day 4, the transcriptomes diverged after in response to PXGL or TSC media, and gradually followed different trajectories toward naive-iPSCs and stabilised TSCs samples (Fig. 4c).

**FIGURE 4.**
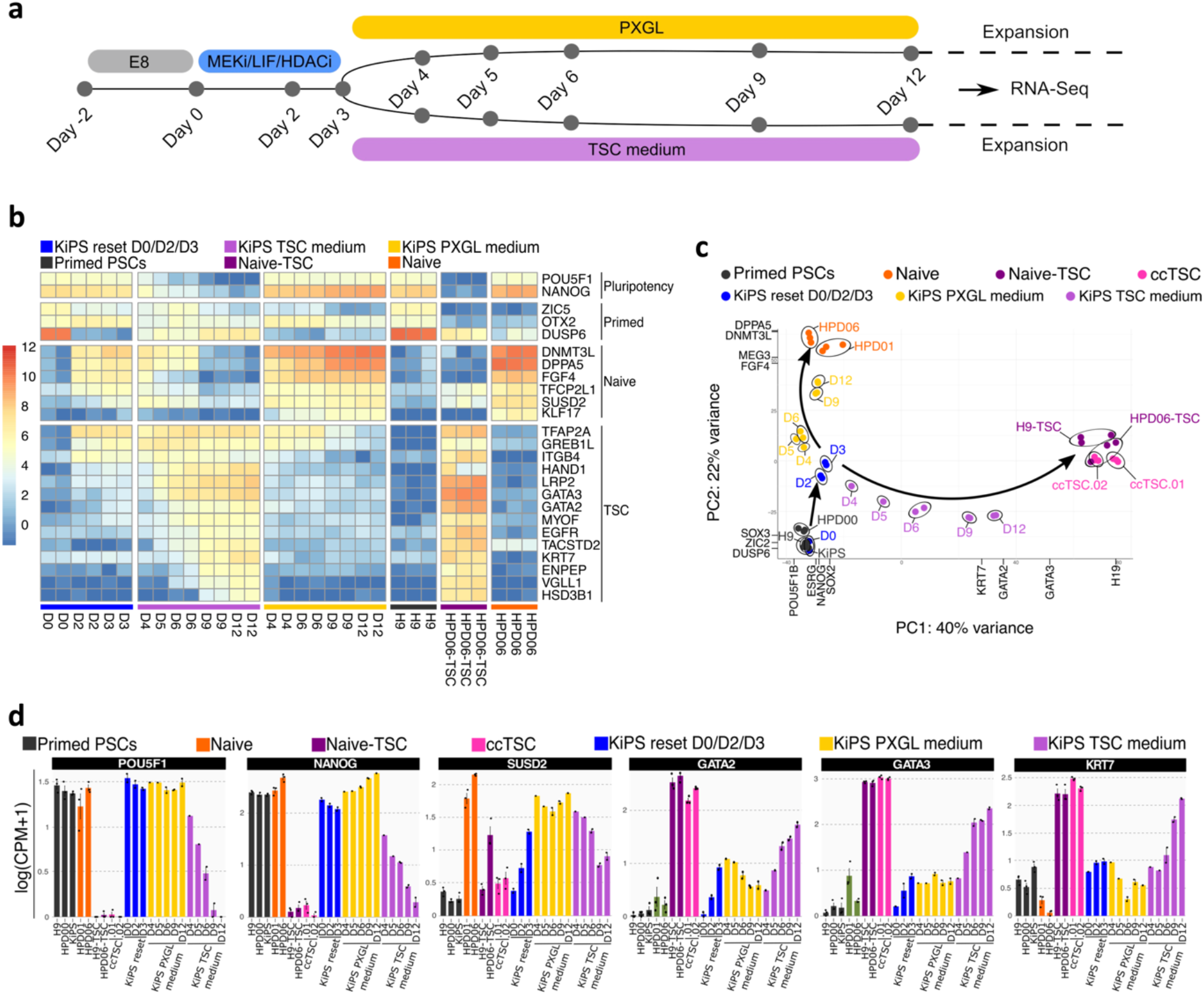
Transcriptional analysis of ccTSCs generation. **a** Experimental scheme of the conversion of conventional PSCs into naive PSCs or TSCs. Time points for RNA-seq sample collection are shown. **b** Heatmap of primed, naive and trophoblast specific genes during the conversion of conventional PSCs into TSCs or naive PSCs, with primed-ESCs (H9), TSCs derived from naive PSCs (HPD06 naive-TSCs), and naive-iPSCs (HPD06). Row-scaled log2(CPM) expression values are shown. Red and blue indicate high and low expression, respectively. Main clusters have been identified as marker genes for general pluripotency, primed-iPSCs, naive-iPSCs and TSCs derived from naive PSCs. **c** Principal component analysis of the time point conversion of conventional PSCs into naive PSCs or TSCs, with primed-PSC (HPD00, H9 and KiPS), naive-iPSCs (HPD06 and HPD01), TSCs derived from naive PSCs (HPD06 naive-TSCs and H9 naive-TSCs), and ccTSCs (ccTSC.01 and ccTSC.02), and performed on the top 5000 most variable genes identified through RNA-seq. Conditions are highlighted in different colours and biological replicates of the same cell line are labelled. **d** Barplots showing the absolute expression as log(CPM+1) of general pluripotency (OCT4 and NANOG), naive (SUSD2), and TSC specific genes (GATA2, GATA3 and KRT7) in the reported conditions highlighted in different colours. Bars indicate the mean ± SEM of independent experiments shown as dots.

Expression of naive-specific markers (e.g. SUSD2 and TFCP2L1) were further consolidated in PXGL, or gradually switched off in TSC medium (Fig. 4b-d). Similarly, the expression of TSC markers, like TFAP2A, was detected early during the chemical resetting, and gradually decreased in PXGL or further increased in TSC medium, together with other markers such as GATA2/3 and KRT7 (Fig. 3b-d). Our data indicate that divergent trajectories toward naive-iPSCs and stabilised TSCs proceed through a shared intermediate state, which can be reached in 3 days of chemical resetting and in which the cell population is highly responsive to differentiation stimuli.

## Discussion

We reported the rapid and efficient conversion of conventional PSCs to TSCs without the need of genetic modifications or prolonged expansion under conditions supporting naive pluripotency. Previous studies reported resetting of conventional PSCs to naive PSCs, followed by differentiation towards TE and to TSCs^4,6,17,19^. These multi-step protocols require the isolation of the cell type of interest by fluorescence-activated cell sorting and long-term expansion, possibly to enrich some rare populations. In particular, a stable naive phenotype is obtained only after isolation and expansion for some passages in naive pluripotency-sustaining conditions. We were thus rather surprised by the rapidity with which we obtained a homogeneous TSCs population with negligible expression of amnion markers from conventional PSCs through histone deacetylase inhibition. Such conversion might occur either directly or via a transient naive pluripotent state. Interestingly, in the case of reprogramming of somatic cells, a trophoblast-like identity was transiently acquired *before* naive pluripotency establishment^35^, raising the possibility that also during chemical resetting cells acquire a transient TE or trophoblast identity on their path towards the naive state. Conversely, recent findings also proposed the co-expression of embryonic and extraembryonic genes as a functional attribute of naive pluripotency, suggesting that the trophoblast identity can be acquired via a transient naive-like state^17^. In our work, we detected co-expression of naive PSCs and TSCs markers at the population levels after 3 days of chemical conversion, indicating either the presence of a mixed cell population or the generation of an intermediate cellular state responsive to different external stimuli. Future single-cell analyses will reveal the actual trajectories followed by cells acquiring TSC and naive PSC identities. The conversion occurs in the presence of HDACi, LIF and the MEK inhibitor. The HDACi and LIF might affect the epigenome of conventional PSCs, by increasing histone acetylation and reducing DNA methylation^44^. It should also be noted that HDAC inhibitors are used as anti-cancer drugs as they induce cell cycle arrest and apoptosis in cancer cell lines. HDAC inhibitors are also used to enhance reprogramming or cell differentiation, but rarely for cell expansion. Conversely, the HDAC inhibitor VPA allows for the robust expansion of TSCs^3^, suggesting that high levels of histone acetylation might be an intrinsic and peculiar feature of human TSCs. In this view, our findings also imply that epigenetic modifications and extensive methylation could be a limiting factor in the competence of conventional PSCs^22–24^ to directly form *bona fide* TSCs.

LIF might promote the conversion by boosting oxidative phosphorylation^45^ or by inducing and maintaining naive pluripotency, as reported in murine cells ^45–47^, although it is still not clear whether LIF is strictly required to sustain naive pluripotency in human cells^8,11,20^.

MEK inhibition is commonly used to stabilise naive pluripotency in human^8,10,12,20,21,48^ and murine cells^22,49^. MEK inhibition has also been shown to promote the transition of naive PSCs towards TE^4,6^ in the absence of other stimuli supporting human naive pluripotency, such as TGF-beta signalling, PKC inhibition and SRC inhibition^8,10–12,20,21,50^. Thus MEK inhibition seems to have a dual role, as stabiliser of human naive pluripotency and inducer of TE identity. This could explain the enhanced plasticity of human naive PSCs.

It will be interesting to test which signalling pathways, epigenetic modifications and metabolic pathways are specifically needed for the generation of naive PSCs or TSCs from conventional PSCs. Our results have practical applications, given that conventional iPSC lines have been generated from patients with placental disorders^39^. It will be extremely easy to convert such iPSCs to TSCs, from which placental organoids or specific placental cell types of interest could be obtained.

## Methods

### Culture of hPSCs

Human primed hiPSCs (KiPS^10^, Keratinocytes induced Pluripotent Stem Cells and HPD00^13^) and hESCs (H9) were cultured in Feeder-free on pre-coated plates with 0.5% growth factor-reduced Matrigel (CORNING 356231) (vol/vol in PBS with MgCl2/CaCl2, Sigma-Aldrich D8662) in E8 medium (made in-house according to^51^) at 37°C, 5% CO_2_, 5% O_2_. Cells were passaged every 3-4 days at a split ratio of 1:8 following dissociation with 0.5 mM EDTA (Invitrogen AM99260G) in PBS without MgCl2/CaCl2 (Sigma-Aldrich D8662), pH8. KiPS line was derived by reprogramming of human keratinocytes^10^ (Invitrogen) with Sendai viruses encoding for OSKM and kindly provided by Austin Smith’s laboratory.

Human naïve ESCs (reset H9, were provided by Austin Smith’s laboratory and were generated via transient expression of NK2 and previously described in^10^) and human naïve iPSCs (HPD06 and HPD01 were previously generated by direct reprogramming from somatic cells and described in^13^) were cultured on mitotically inactivated mouse embryonic fibroblasts (MEFs; DR4 ATCC) in PXGL medium^36^ or in RSeT medium (Stem Cell Technologies 05969). The PXGL medium was prepared as follows: N2B27 (DMEM/F12 [Gibco 11320-074], and Neurobasal in 1:1 ratio [Gibco 21103-049], with 1:200 N2 Supplement [Gibco 17502-048], and 1:100 B27 Supplement [Gibco 17504-044], 2 mM L-glutamine [Gibco 25030-024], 0.1 mM 2-mercaptoethanol [Sigma-Aldrich M3148]) supplemented with 1µM PD0325901 (Axon Medchem), 2 μM XAV939 (Axon Medchem), 2 μM Gö6983 (Axon Medchem) and 10 ng/ml human LIF (produced in-house). Human naive PSCs were passaged as single cells every 4 days at split ratio 1:3 or 1:4 following dissociation with TrypLE (Gibco 12563-029) for 10 minutes (min) at room temperature (RT). The ROCK inhibitor (Y27632, Axon Medchem 1683) was added in the naïve medium only for 24h after passaging.

All cell lines were mycoplasma-negative (Mycoalert, Lonza).

### Culture of hTSCs

hTSCs were cultured as previously described^3^. Briefly, a 12-well plate was coated with 5 μg/mL Collagen IV (Corning, 354233) at 37°C overnight or on mitotically inactivated mouse embryonic fibroblasts. Cells were cultured in 0.8 mL TS medium [DMEM/F12 supplemented with 0.1 mM 2-mercaptoethanol, 0.2% FBS, 0.5% Penicillin-Streptomycin, 0.3%, BSA (Gibco 15260-037), 1% ITS-X (Gibco, 51500), 1.5 μg/ml L-ascorbic acid (Sigma A4544), 50 ng/ml EGF (peprotech AF-100-15), 2 μM CHIR99021 (Axon Medchem cat. nos 1386 and 1408), 0.5 μM A83-01 (Axon Medchem 1421), 1 μM SB431542 (Axon Medchem 1661), 0.8 mM VPA (HDACi, Sigma, P4543), and 5 μM Y-27632] and in 5% CO2 and 5% O2. Media were changed every 2 days, and cells were passed using TrypLE Express every 3-4 days at a ratio of 1:8. Unless otherwise specified, hTSCs between passage 5 and 20 were used for experiments. All cell lines were mycoplasma-negative (Mycoalert, Lonza).

### Culture of BAP-treated primed pluripotent stem cells

KiPS were dissociated into single cells with TrypLE. The cells were cultured on pre-coated plates with 0.5% growth factor-reduced Matrigel at a density of 2×10^4^ cells/cm^2^ with MEF-conditioned medium supplemented with 10 ng/ml BMP4 (Peprotech, 120-05ET), 1µM A83-01, 0.1µM PD173074 (Axon, 1673), and 10µM Y-27632, as described previously^4^. DMEM/Ham’s F-12 medium containing 0.1 mM 2ME, 1% ITS-X supplement, 1% NEAA, 2 mM L-glutamine, and 20% KSR was cultured on MEF for 24 hours, and the supernatant was collected and used as MEF-conditioned medium. The medium was changed daily.

### Chemical resetting

For the chemical resetting from primed to naive PSC, KiPS were seeded at 10000 cells/cm^2^ on mitotically inactivated MEFs in E8 medium with 10µM ROCKi (added only for 24 hours). Two days after plating (day 0), medium was changed to PD03/LIF/HDACi [N2B27 with 1µM PD0325901, 10ng/ml human LIF and 1mM VPA (HDACi)]. Following 3 days in PD03/LIF/HDACi, medium was changed to PXGL for a further 10-11 days before passing in PXGL medium on MEFs (see Fig. 1a). For the chemical resetting from primed to ccTSCs, KiPS were seeded at 10000 cells/cm^2^ on mitotically inactivated MEFs in E8 medium. Two days later (day 0) medium was changed to PD03/LIF/HDACi. Following 3 days in PD03/LIF/HDACi, medium was changed to PXGL for a further 4 days and then switched with TSC medium in the case of ccTSC.01 (see Fig. 2f) or directly to TSC medium in the case of ccTSC.02 (see Fig. 3a). After resetting, ccTSCs were passaged as TSCs (see above).

### Conversion of human naive PSCs into TSCs

Naive hPSCs did a pre-treatment on MEF with TS medium for 24h, whereupon naive hPSCs were single-cell dissociated by TrypLE Express, and 0.25–0.5 × 10^6^ cells were seeded on a 12-well plate pre-coated with 5 μg/mL Collagen IV or on mitotically inactivated mouse embryonic fibroblasts and cultured in 0.8 mL TS medium. Cells were cultured in 5% CO2 and 5% O2, media was changed every 2 days, and passaged upon 80–100% confluency at a ratio of 1:2 to 1:4. During derivation from naive hPSCs, the cells grew slowly during the initial few passages. Between passage 5 and 10, highly proliferative hTSCs emerged.

### Differentiation of STB

Differentiation of hTSCs into terminal cell types was performed as previously described^3^, with minor modifications. Prior to differentiation, hTSCs were grown to about 80% confluency, and then single-cell dissociated using TrypLE Express. For STB differentiation, 12-well plates were coated with 2.5 μg/mL Collagen IV overnight. 1 × 10^5^ hTSCs were seeded per well in 0.8 mL STB medium [DMEM/F12 supplemented with 0.1 mM β-mercaptoethanol, 0.5% penicillin-streptomycin, 0.3% BSA, 1% ITS-X, 2.5 μM Y-27632, 2 μM Forskolin (Sigma-Aldrich, F3917), and 4% KSR]. The media was changed at day 3, and at day 6 the cells were ready for downstream analysis.

### Trophoblast organoids

To obtain trophoblast Organoids we used the protocol previously described in^42^. TSCs were single-cell dissociated by TrypLE Express and approximately 50K cells were resuspended in Matrigel (Corning 356231) on ice. Drops (25 µl) were plated per well into a 24-well culture plate, set at 37 °C for 10 min and overlaid with 400 µl trophoblast organoid medium (TOM, previously described in^42^). Cultures were maintained in 5% CO2 and 5% O2 in a humidified incubator at 37 °C. Medium was replaced every 2– 3 days. Small organoid clusters became visible by around day 7 and were passaged when at least 50% had reached a diameter of 200–300 µm (usually between days 7 and 10). Mechanical disruption was achieved by pipetting the drops of Matrigel with TrypLE Express to obtain a single cell suspension, after that spin dawn at 4°C and resuspend the pellet directly in Matrigel at the correct proportion. Frozen stocks of organoids were made in 70% TOM, 20% FBS and 10% DMSO freezing medium and stored in liquid nitrogen.

### Immunofluorescence

Immunofluorescence analysis was performed on 1% Matrigel-coated glass coverslip in wells. Cells were fixed in 4% Formaldehyde (Sigma-Aldrich 78775) in PBS for 10 min at RT, washed in PBS, permeabilized for 1 hour in PBS + 0.3% Triton X-100 (PBST) and in the case of TSCs we permeabilized only for 10 min in PBST at RT in order to avoid cell detachment, and blocked in PBST + 5% of Horse serum (ThermoFisher 16050-122) for 5 hours at RT. Cells were incubated overnight at 4°C with primary antibodies (See Supplementary Table 1) in PBST+ 3% of Horse serum. After washing with PBS, cells were incubated with secondary antibodies (Alexa, Life Technologies) (Supplementary Table 1) for 45 min at RT. Nuclei were stained with either DAPI (4′,6-diamidino-2-phenylindole, Sigma-Aldrich F6057).

Images were acquired with a Zeiss LSN700, a Leica SP5 or a Leica SP8 confocal microscope using ZEN 2012 or LeicaLeica TCS SP5 LAS AF (v2.7.3.9723) software respectively.

### Quantitative PCR

Total RNA was isolated using Total RNA Purification Kit (Norgen Biotek 37500), and complementary DNA (cDNA) was made from 500 ng using M-MLV Reverse Transcriptase (Invitrogen 28025-013) and dN6 primers. For real-time PCR SYBR Green Master mix (Bioline BIO-94020) was used. Primers are detailed in Supplementary Table 2. Three technical replicates were carried out for all quantitative PCR. GAPDH was used as an endogenous control to normalise expression. qPCR data were acquired with QuantStudio™ 6&7 Flex Software 1.0.

### RNA sequencing and analysis

Quant Seq 3’ mRNA-seq Library Prep kit (Lexogen) is used for library construction. Library quantification is performed by fluorometer (Qubit) and bioanalyzer (Agilent). Sequencing is performed on NextSeq500 ILLUMINA instruments to produce 5 million reads (75bp SE) for the sample.

For the analysis of imprinted genes and TSCs markers, the reads were trimmed using BBDuk (BBMap v. 37.87), with parameters indicated in the Lexogen data analysis protocol. After trimming, reads were aligned to the Homo sapiens genome (GRCm38.p13) using STAR (v. 2.7.6a). The gene expression levels were quantified using featureCounts (v. 2.0.1). Genes were sorted removing those that had a total number of counts below 10 in at least 3 samples. All RNA-seq analyses were carried out in the R environment (v. 4.0.0) with Bioconductor (v. 3.7). We computed differential expression analysis using the DESeq2 R package (v. 1.28.1)^52^. Transcripts with absolute value of log2[FC] > 1 and an adjusted p-value < 0.05 (Benjamini–Hochberg adjustment) were considered significant and defined as differentially expressed for the comparison in the analysis. Heatmaps were made using counts-per-million (CPM) values with the pheatmap function from the pheatmap R package (v.1.0.12) on differentially expressed genes or selected markers. Volcano plots were computed with log2[FC] and − log10[adjusted p-value] from DESeq2 differential expression analysis output using the ggscatter function from the ggpubr R package (v. 0.4.0). Barplots were made using CPM values with the ggbarplot function from the ggpubr R package.

For the time point analysis during resetting and the following stabilisation in PXGL or TSC medium, transcript quantification was performed from raw reads using Salmon (v1.6.0)^53^ on transcripts defined in Ensembl 105. Gene expression levels were estimated with tximport R package (v1.20.0)^54^ and differential expression analysis was computed using the DESeq2 R package (v. 1.28.1)^52^. Transcripts with absolute value of log2[FC] ≥ 3 and an adjusted p-value < 0.05 (Benjamini–Hochberg adjustment) were considered significant. Principal component analysis was performed on variance stabilised data (vst function from DESeq2 R package v 1.32.0^52^) using prcomp function on the top 5000 most variable genes. Heatmaps were performed using the pheatmap function (pheatmap R package v1.0.12) on log2 count per million (CPM) data of selected markers. All analysis except salmon were performed in R version 4.1.1.

## Statistics and reproducibility

For each dataset, sample size *n* refers to the number of independent experiments or biological replicates, shown as dots, as stated in the figure legends. R software (v4.0.0) was used for statistical analysis. All error bars indicate the standard error of the mean (SEM). All qPCR experiments were performed with three technical replicates.

## Data availability

RNA-seq data for this study have been deposited in the gene Expression Omnibus (Geo) database under the accession code GSE184562.

**Supplementary Table 1.**

| Antibody | Dilution | Product code | Company |
| --- | --- | --- | --- |
| OCT4 | 1:300 | sc-5279 | Santa Cruz |
| KLF17 | 1:250 | HPA024629 | Atlas |
| KRT7 | 1:100 | AB183344 | Abcam |
| GATA3 | 1:100 | AF2605 | R&D |
| Donkey anti-Goat IgG (H+L)<br>Secondary Antibody Alexa<br>Fluor 488 | 1:500 | A-11055 | ThermoFisher |
| Donkey anti-Mouse IgG (H+L)<br>Secondary Antibody Alexa Fluor 568 | 1:500 | A-10037 | ThermoFisher |
| Donkey anti-Rabbit IgG (H+L)<br>Secondary Antibody Alexa Fluor 647 | 1:500 | A-31573 | ThermoFisher |

**Supplementary Table 2.**
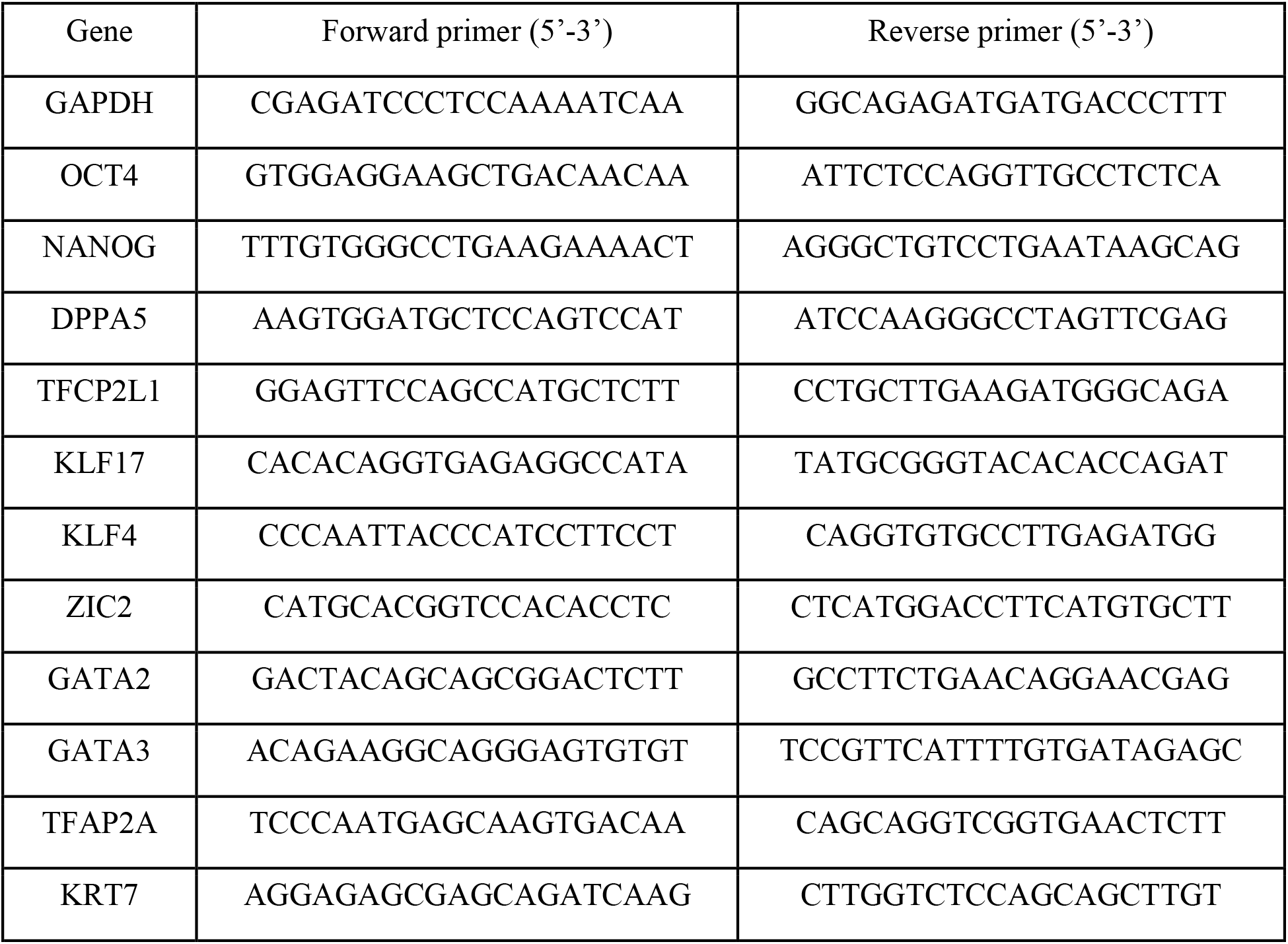
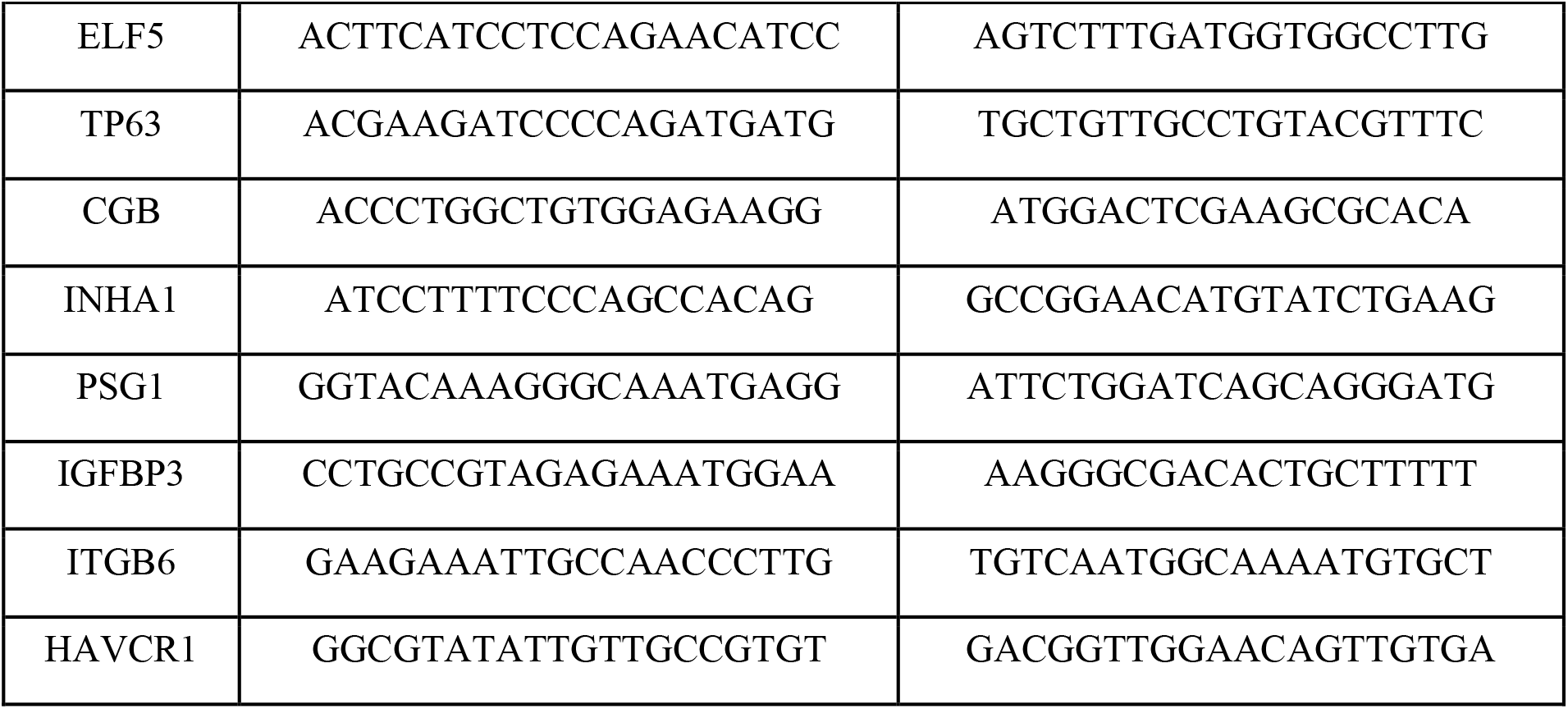

**SUPPLEMENTARY FIGURE 1.**
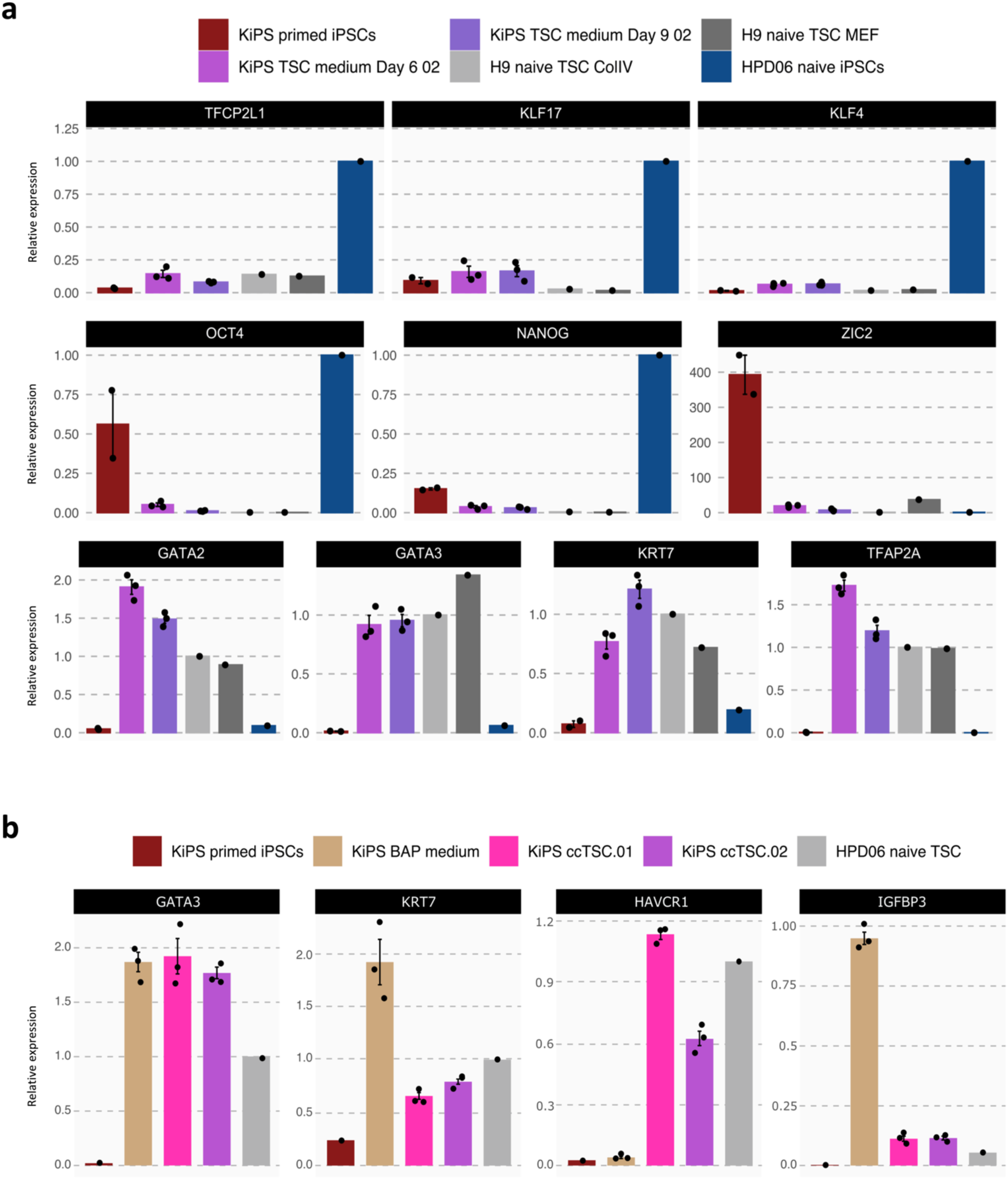
**a** Gene expression analysis by RT–qPCR of KiPS primed iPSCs, KiPS in TSC medium at day 6 and 9, H9 naive-TSC cultured on ColIV or MEF and HPD06 naive iPSCs. Top and middle: naive, primed and general pluripotency markers, expression was normalised to HPD06 naive iPSCs; bottom: trophoblast markers, expression was normalised to H9 naive-TSC. Bars indicate the mean ± SEM of biological replicates shown as dots. **b** Gene expression analysis by qPCR of KiPS primed iPSCs, KiPS in BAP medium, KiPS ccTSC.01, KiPS ccTSC.02, and HPD06 naive TSC. For trophoblast markers the expression was normalised to HPD06 naive TSC, for amnion markers the expression was normalised to the mean of KiPS in BAP medium samples. Bars indicate the mean ± SEM of biological replicates shown as dots.

## Acknowledgements

The authors thank the other members of the Martello Laboratory for discussions, suggestions and technical support. G.M.’s Laboratory is supported by grants from the Giovanni Armenise–Harvard Foundation, the Telethon Foundation (TCP13013) and an ERC Starting Grant (MetEpiStem).

## Author information

Irene Zorzan Present address: Epigenetics Programme, Babraham Institute, CB22 3AT, Cambridge, UK.

These authors contributed equally: Irene Zorzan and Riccardo Massimiliano Betto

## Contributions

G.M. designed the study; I.Z. and A.D. performed chemical resetting; R.M.B and I.Z. established and characterised TSC lines from naive PSCs; R.M.B. and I.Z. performed TSC differentiation and organoid formation assays; M.A., G.R. and P.M. performed bioinformatic analyses; I.Z, R.M.B and A.D performed immunostaining and qPCR; M.A., I.Z., R.M.B, G.R. and A.D. prepared the figures; G.M. wrote the manuscript with input from all authors; G.M. and C.R. supervised the study and provided fundings.

## Corresponding authors

Correspondence to Graziano Martello.

